# Feature similarity gradients detect alterations in the neonatal cortex associated with preterm birth

**DOI:** 10.1101/2022.09.15.508133

**Authors:** Paola Galdi, Manuel Blesa Cabez, Christine Farrugia, Kadi Vaher, Logan ZJ Williams, Gemma Sullivan, David Q Stoye, Alan J Quigley, Antonios Makropoulos, Michael J Thrippleton, Mark E Bastin, Hilary Richardson, Heather Whalley, A David Edwards, Claude J Bajada, Emma C Robinson, James P Boardman

**Affiliations:** MRC Centre for Reproductive Health, University of Edinburgh, UK; Department of Physiology and Biochemistry, Faculty of Medicine and Surgery, University of Malta, Malta; Centre for the Developing Brain, King’s College London, UK; School of Biomedical Engineering and Imaging Science, King’s College London, UK; Centre for Clinical Brain Sciences, University of Edinburgh, UK; Royal Hospital for Children & Young People, Edinburgh, UK; School of Philosophy, Psychology and Language Sciences, University of Edinburgh, UK; Centre for Genomic and Experimental Medicine, University of Edinburgh, UK; MRC Centre for Neurodevelopmental Disorders, King’s College London, UK; University of Malta Magnetic Resonance Imaging Platform (UMRI)

## Abstract

The early life environment programmes cortical architecture and cognition across the life course. A measure of cortical organisation that integrates information from multi-modal MRI and is unbound by arbitrary parcellations has proven elusive, which hampers efforts to uncover the perinatal origins of cortical health. Here, we use the Vogt-Bailey index to provide a fine-grained description of regional homogeneities and sharp variations in cortical microstructure based on feature gradients, and we investigate the impact of being born preterm on cortical development. Preterm infants have a homogeneous microstructure in temporal and occipital lobes, and the medial parietal, cingulate, and frontal cortices, compared with term infants. These observations replicated across two independent datasets and were robust to differences that remain in the data after matching samples and alignment of processing and quality control strategies. We conclude that cortical microstructural architecture is altered in preterm infants in a spatially distributed rather than localised fashion.

## Introduction

Parcellation of the brain into spatially contiguous and non-overlapping regions with a locally homogeneous profile is a useful abstraction to study cerebral function and organisation. However, there is no consensus on the optimal delineation of brain parcels, whether they are defined cytoarchitectonically, on the basis of function, or on a combination of features (***Bohland et al., 2009***; ***Cloutman and Lambon Ralph, 2012***; ***Arslan et al., 2018***; ***Eickhoff et al., 2018***; ***Glasser et al., 2016***). Individual variability in brain shape and organisation further complicates the definition of an universal brain map, as reference templates might not adapt well to the signal of individual acquisitions (***Eickhoff et al., 2018***). One of the main challenges in defining parcels is the arbitrary assignment of sharp boundaries between regions that may only show graded differences in local structure or function. ***Glasser et al. (2016***) attempted to overcome this issue by proposing a method to detect boundaries between parcels that observed gradients (i.e. spatial transitions) in multimodal brain maps describing different brain features: myelin-sensitive features (derived from T1/T2 contrasts), cortical thickness, cortical folding, and functional activity patterns and connectivity.

Feature similarity gradients provide a unified approach to study graded differences in brain structure and function (***Bajada et al., 2017, 2020***; ***Bernhardt et al., 2022***). This technique has been applied to study the cortical organisation of the adult brain, revealing the existence of a global gradient with primary sensory and motor regions at one end of the spectrum and heteromodal regions at the opposite end (***Huntenburg et al., 2018***; ***Margulies et al., 2016***). This gradient is already detectable during adolescence (***Dong et al., 2021***). Closely after birth and through childhood the primary axis of cortical organisation is anchored within the unimodal cortex, between somatosensory, motor and visual regions, whereas higher order functional systems develop later (***Larivière et al., 2020***; ***Dong et al., 2021***).

The cortex undergoes dramatic changes in the last trimester of pregnancy and early postnatal life as a result of concurrent cellular and molecular processes including synaptogenesis, axonal growth, dendritic arborization and myelination (***Ouyang et al., 2018***; ***Volpe, 2019***; ***Kostović et al., 2019***). Disruption of normal gestation during this period of rapid growth can have long lasting consequences on neurodevelopment (***Zhang et al., 2015***; ***Kersbergen et al., 2016***; ***Fleiss et al., 2020***; ***Kline et al., 2020***). Imaging of the preterm brain at term-equivalent age has characterised dysmaturation following preterm birth as a global phenomenon (i.e. encompassing the whole brain), with generalised dysconnectivity and atypical topology of developing neural networks, increased water diffusivity, altered white matter microstructure and reduced brain volume and cortical surface area (***Telford et al., 2017***; ***Batalle et al., 2018***; ***Boardman and Counsell, 2019***; ***Blesa et al., 2021***; ***Vaher et al., 2022***). Motivated by the search of neural antecedents of cognitive impairment observed in some preterm born individuals, studies focusing on the neonatal cortex have demonstrated that prematurity impacts cortical growth and microstructural development in a dose-dependent fashion (***Kapellou et al., 2006***; ***Ball et al., 2013b***) and that preterm birth alters cortical folding (***Shimony et al., 2016***; ***Neil and Smyser, 2018***; ***Dubois et al., 2019***), thalamo-cortical connectivity (***Boardman et al., 2006***; ***Ball et al., 2013a***), and different regional MRI metrics with varying spatial distribution, including water diffusion measures, markers of myelination and cortical morphology metrics (***Bouyssi-Kobar et al., 2018***; ***Ouyang et al., 2019***; ***Ball et al., 2020***; ***Dimitrova et al., 2021***). Additionally, using principal component analysis of multimodal MRI metrics, ***Ball et al. (2020***) found that prematurity leads to alterations along the principal axis of variation in cortical microstructure.

Here we propose a framework to study feature similarity gradients in the microstructural organisation of the neonatal cortex based on six microstructural metrics: fractional anisotropy, mean, axial and radial diffusivities, neural density index and orientation dispersion index. We apply it to study the impact of being born preterm on cortical development, and we test replicability in two independent datasets: the Theirworld Edinburgh Birth Cohort (TEBC, ***Boardman et al. (2020***)) and the developing Human Connectome Project (dHCP, ***Edwards et al. (2022***)). The approach is based on the Vogt-Bailey index (VB), previously proposed by ***Bajada et al. (2020***). By integrating information from multiple features computed at each point of the cortical mantle, this index measures the extent of discontinuity in cortical intra-areal relationships, summarising in a single metric the information contained in a feature similarity gradient. Compared to previous approaches based on parcels or whole-brain gradients, this method works on a more local scale, providing a fine-grained description of regional homogeneities and sharp variations in cortical properties. An additional advantage of this method is that local homogeneity is computed in subject native space, obviating the need of registering volumes to a common template and allowing for greater anatomical specificity.

## Results

In the TEBC sample, 254 infants had multimodal MRI data suitable for the proposed analysis. Quality control was performed separately for the left and right hemispheres. After visual inspection of the cortical surfaces, 221 subjects (147 preterm and 72 term-born infants) were included in either the left or right hemisphere analysis, or both (Table 1). Of the preterm infants, 36 had bronchopulmonary dysplasia (BPD), 8 had necrotising enterocolitis (NEC) and 6 required treatment for retinopathy of prematurity (ROP). After quality control, the dHCP matching sample (matched on age at scan, age at birth and sex distribution) contained 43 preterm and 84 term-born infants (Table 2). None of the preterm infants in the dHCP sample had NEC or ROP; no information was recorded regarding BPD. In both datasets, there were no differences in in-scanner relative motion estimates of volume-to-volume displacement between the preterm and term groups (Table 1). The distributions of gestational age at birth and age at scan for the two datasets are shown in Figure 1. Table 3 reports the sample size for the preterm and term groups for each experiment. There were no statistically significant differences in post-menstrual age (PMA) distributions between the left and right hemisphere samples, or between any individual hemisphere and the whole sample, in either dataset (Wilcoxon rank-sum test, p-values > 0.43).

**Table 1.** Participant characteristics for the TEBC by group. Mean, minimum and maximum values are reported for continuous variables and ratios for binary variables. The last column reports the p-values of the group differences computed with the Wilcoxon rank-sum test for continuous variables and with the chi-squared test for binary variables.

|  | preterm (N=147) | term (N=72) | preterm vs. term |
| --- | --- | --- | --- |
| PMA at birth (weeks) | 29.48 (22.14-32.86) | 39.47 (36.43-42.00) | p = 2.96e-33 |
| Birth weight (grams) | 1318 (370-2510) | 3462 (2410-4560) | p = 3.14e-33 |
| PMA at scan (weeks) | 40.71 (36.57-45.86) | 41.79 (38.29-43.86) | p = 4.23e-9 |
| M:F ratio | 85:62 | 39:33 | p = 4.27e-19 |
| In-scanner motion (mm) | 0.24 (0.10-1.04) | 0.22 (0.12-0.71) | p = 0.4449 |

PMA = Postmenstrual age, M = male, F = female.

**Table 2.** Participant characteristics for the dHCP by group. Mean, minimum and maximum values are reported for continuous variables and ratios for binary variables. The last column reports the p-values of the group differences computed with the Wilcoxon rank-sum test for continuous variables and with the chi-squared test for binary variables.

|  | preterm (N=44) | term (N=89) | preterm vs. term |
| --- | --- | --- | --- |
| PMA at birth (weeks) | 28.98 (23-32.71) | 39.03 (37.14-41.86) | p = 3.51e-20 |
| Birth weight (grams) | 1178 (450-1960) | 3184 (2060-4300) | p = 3.63e-20 |
| PMA at scan (weeks) | 40.86 (38.14-44.29) | 41.07 (37.57-44.71) | p = 0.5838 |
| M:F ratio | 25:18 | 47:37 | p = 7.53e-06 |
| In-scanner motion (mm) | 2.04 (0.75-6.05) | 1.72 (0.65-3.64) | p = 0.1277 |

PMA = Postmenstrual age, M = male, F = female.

**Table 3.** Sample size (N) for each experiment, group and hemisphere.

|  | left hemisphere |  | right hemisphere |  |
| --- | --- | --- | --- | --- |
|  | preterm | term | preterm | term |
| TEBC | 103 | 57 | 132 | 55 |
| dHCP | 37 | 67 | 39 | 70 |

**Figure 1.**
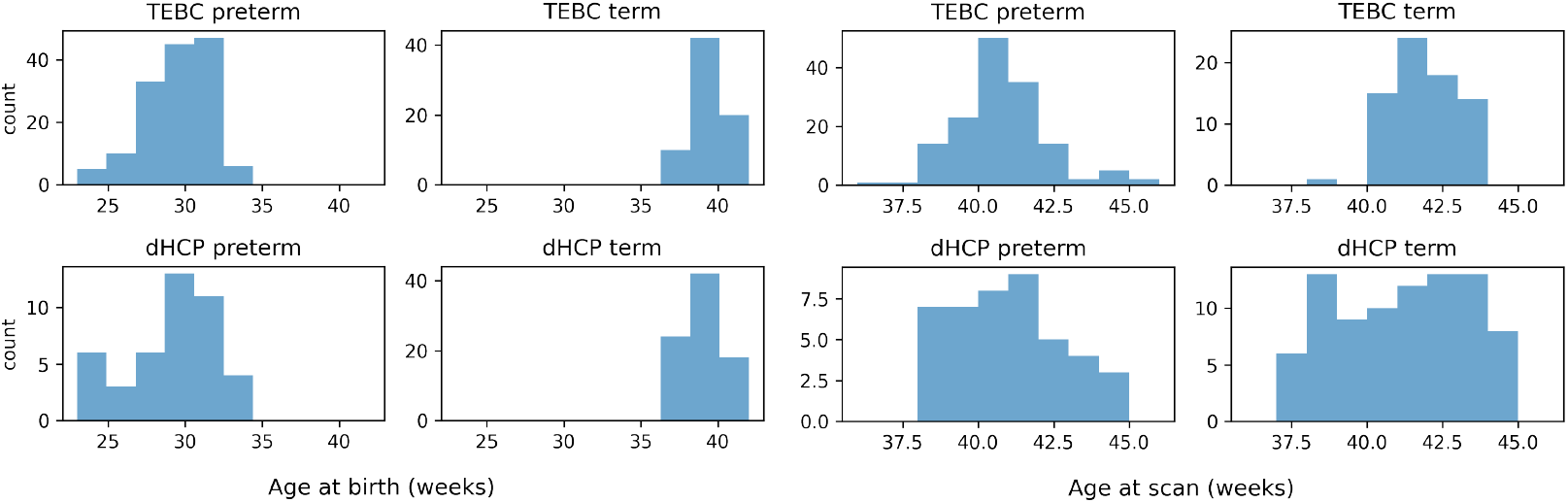
Distribution of postmenstrual age in weeks at birth (left) and at scan (right) for the TEBC and dHCP datasets.

For each subject, VB maps were generated from six microstructural metrics (fractional anisotropy, mean, axial and radial diffusivities, neural density index and orientation dispersion index). Figure 2 shows two exemplar VB maps for one preterm and one term subject.

**Figure 2.**
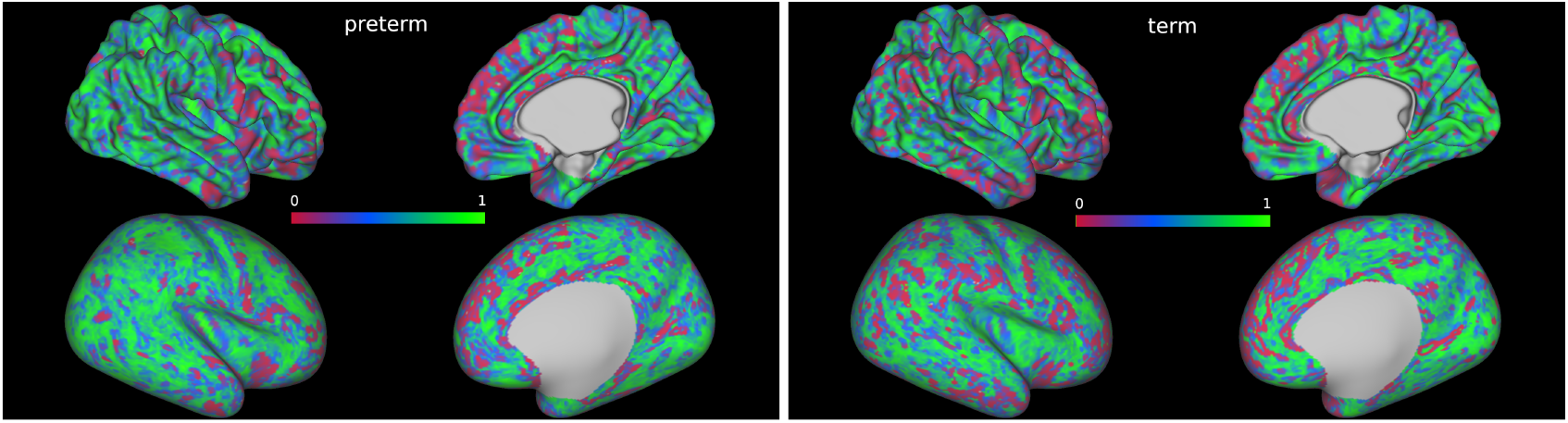
Vogt-Bailey index maps of the right hemisphere of two exemplar subjects, one preterm (left) and one term (right), displayed on midthickness (top row) and inflated (bottom row) surfaces registered to the 40-week dHCP template. Values close to zero indicate local discontinuity in the measured microstructural properties, while values close to 1 indicate a high local homogeneity.

### The impact of preterm birth on cortical organisation

In both datasets, when comparing preterm and term infants controlling for age at scan and sex, we found more homogeneous cortical microstructure (higher VB index) in the preterm group compared to the term group in widespread areas of the cortex including the right temporal lobe, occipital lobe, medial parietal cortex, right cingulate cortex and left frontal cortex. Figure 3 reports the standardised t-maps of the preterm vs. term comparisons for the two datasets, while the p-values maps in Figures 4 and 5 highlight regions where the VB index is significantly higher in the preterm group or in the term group, respectively. In the TEBC dataset only, the VB index is lower (decreased homogeneity) in the preterm group compared to term controls in the medial orbitofrontal cortex. There were no areas of lower VB index in the preterm group in the dHCP dataset.

**Figure 3.**
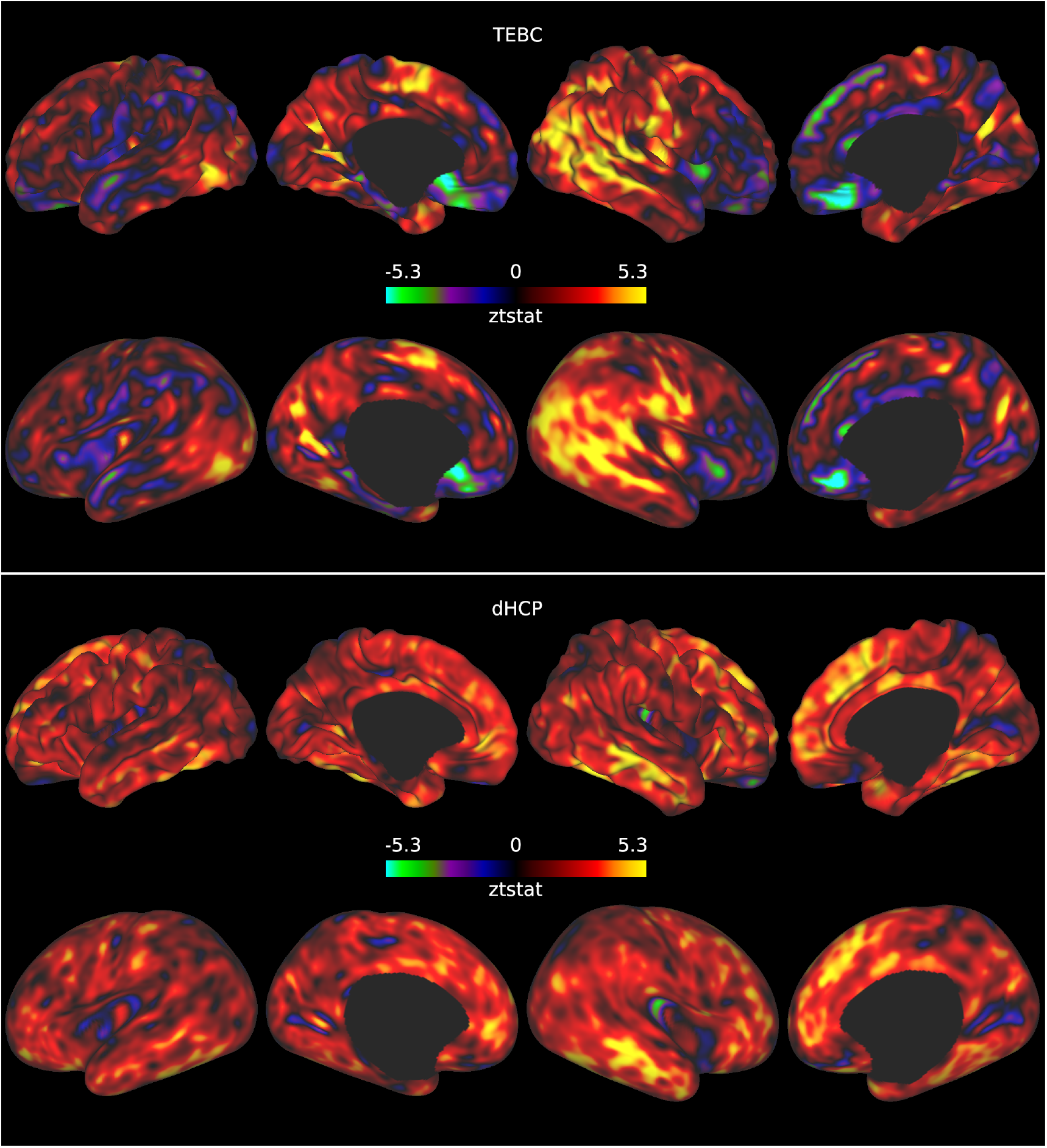
Standardised t-statistic maps resulting from the preterm vs. term comparison for the TEBC sample (top panel) and the dHCP sample (bottom panel). Positive values in the maps indicate regions where the Vogt-Bailey index was higher in the preterm group, and negative values regions where the Vogt-Bailey index was higher in the term group.

**Figure 4.**
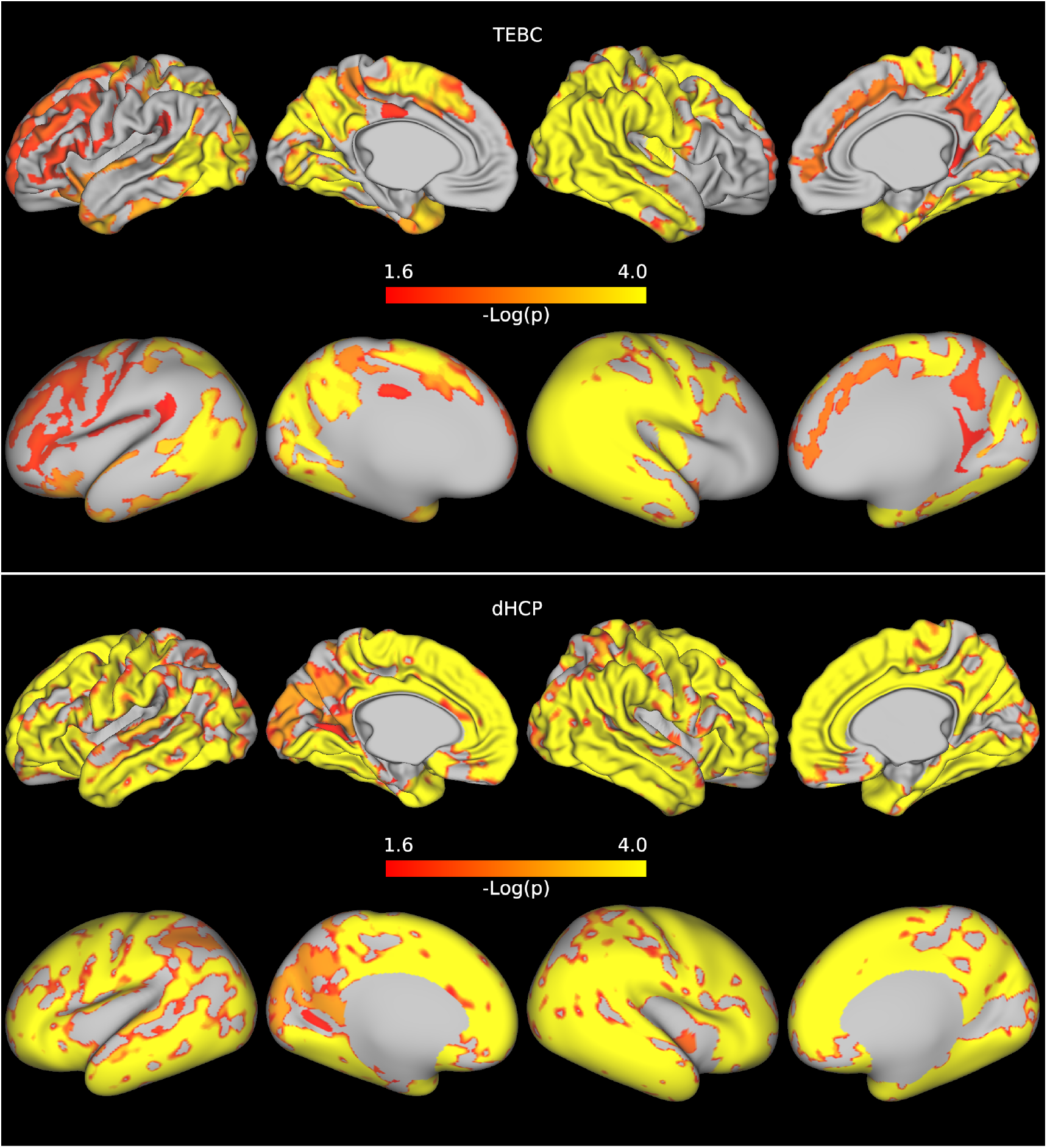
Results of the preterm vs. term comparison for the TEBC sample (top panel) and the dHCP sample (bottom panel). Highlighted areas indicate regions where the Vogt-Bailey index was higher in the preterm group at an alpha level of 0.05 after Šidák correction. The colour bar reports negative log10 p-values.

**Figure 5.**
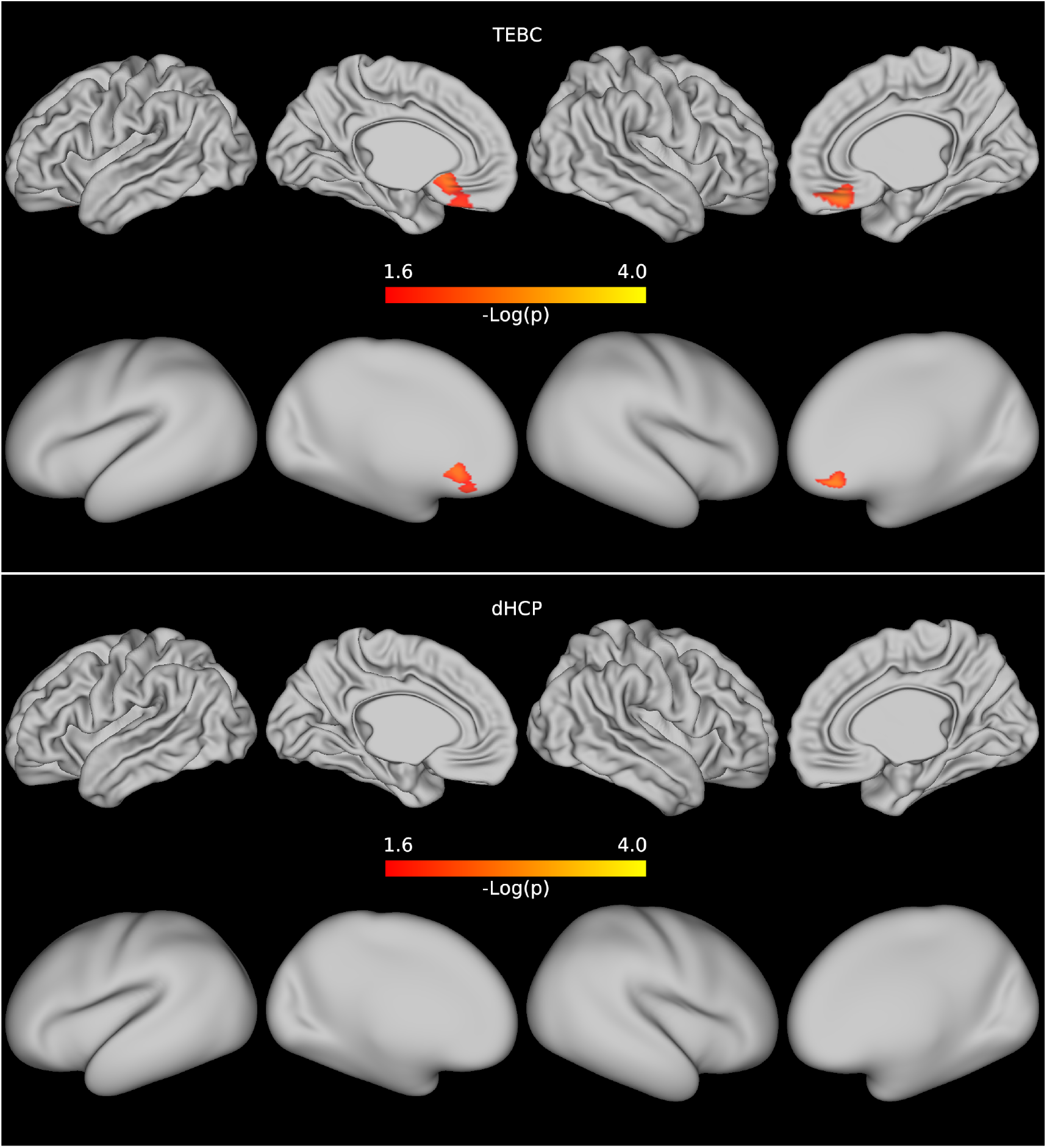
Results of the preterm vs. term comparison for the TEBC sample (top row) and the dHCP sample (bottom row). Highlighted areas indicate regions where the Vogt-Bailey index was higher in the term group at an alpha level of 0.05 after Šidák correction. The colour bar reports negative log10 p-values.

### Replicability across datasets

We tested the replicability of our findings in two datasets, and we aimed to make results as comparable as possible by building matching samples and aligning the structural preprocessing and quality control strategies between the two datasets. However, some discrepancies remain, such as scanner differences, input resolution and variation in acquisition protocols; these had inevitable consequences on the down-stream processing, that are reflected in differences in cortical thickness (Appendix 1) and single-metric maps between the two datasets (Appendix 2) that were spread throughout the cortex.

With the exception of the temporal and occipital lobes, throughout the cortex the VB index was on average higher in the TEBC dataset, with the strongest effect size measured in the medial frontal lobe and right insular cortex (Figures 6-8). There was overlap in between-dataset differences in cortical thickness and between-dataset differences in VB maps in the temporal lobe and insular cortex and in medial parts of the occipital lobe and the frontal gyrus, but not in the rest of the frontal cortex. Single-metric difference maps also showed overlap with the cortical thickness differences but with variable patterns.

**Figure 6.**
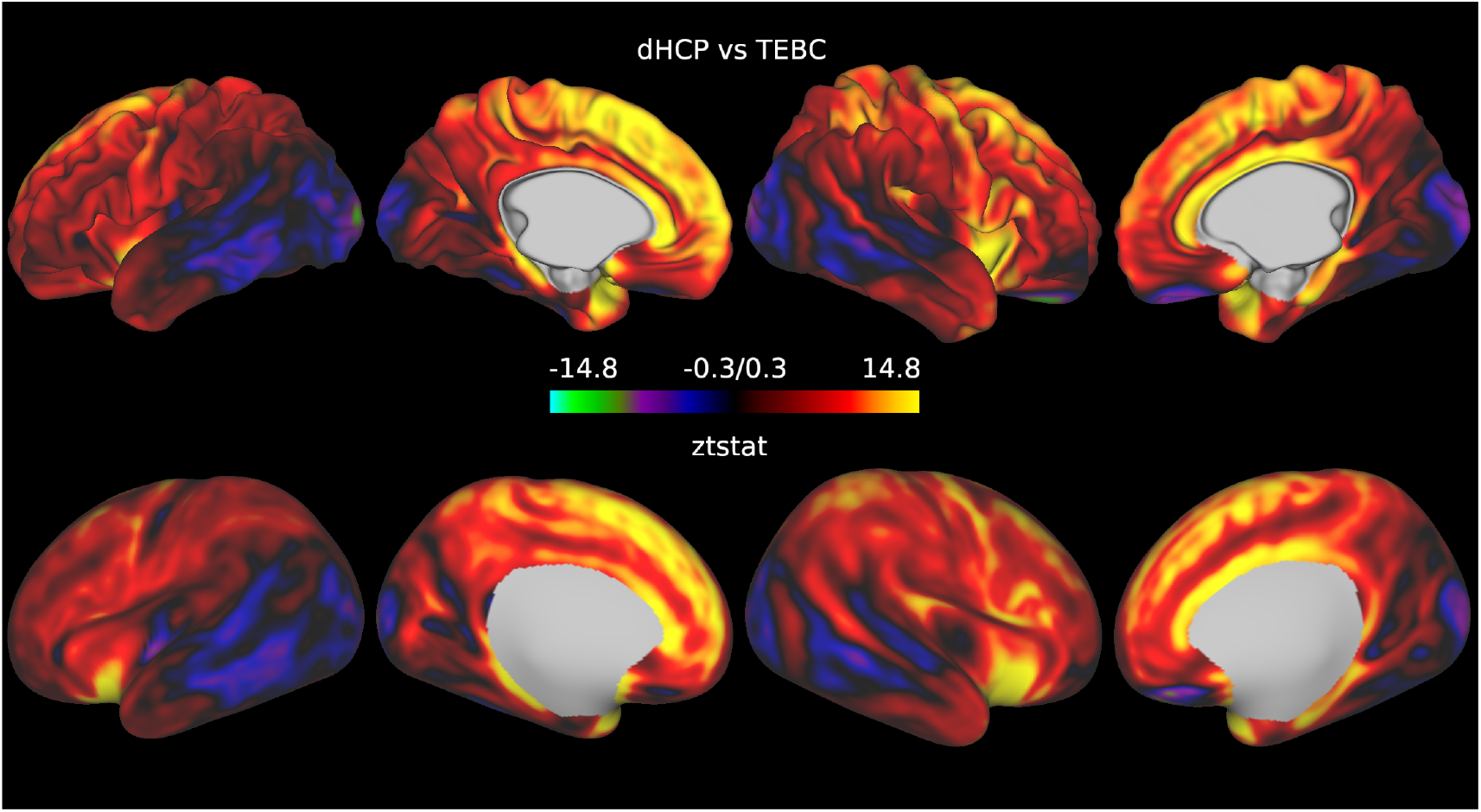
Standardised t-statistic maps showing the results of the comparison of Vogt-Bailey maps between dataset after controlling for prematurity, age at scan and sex. Positive values in the maps indicate regions where the Vogt-Bailey index was higher in the TEBC sample, and negative values regions where the Vogt-Bailey index was higher in the dHCP sample.

**Figure 7.**
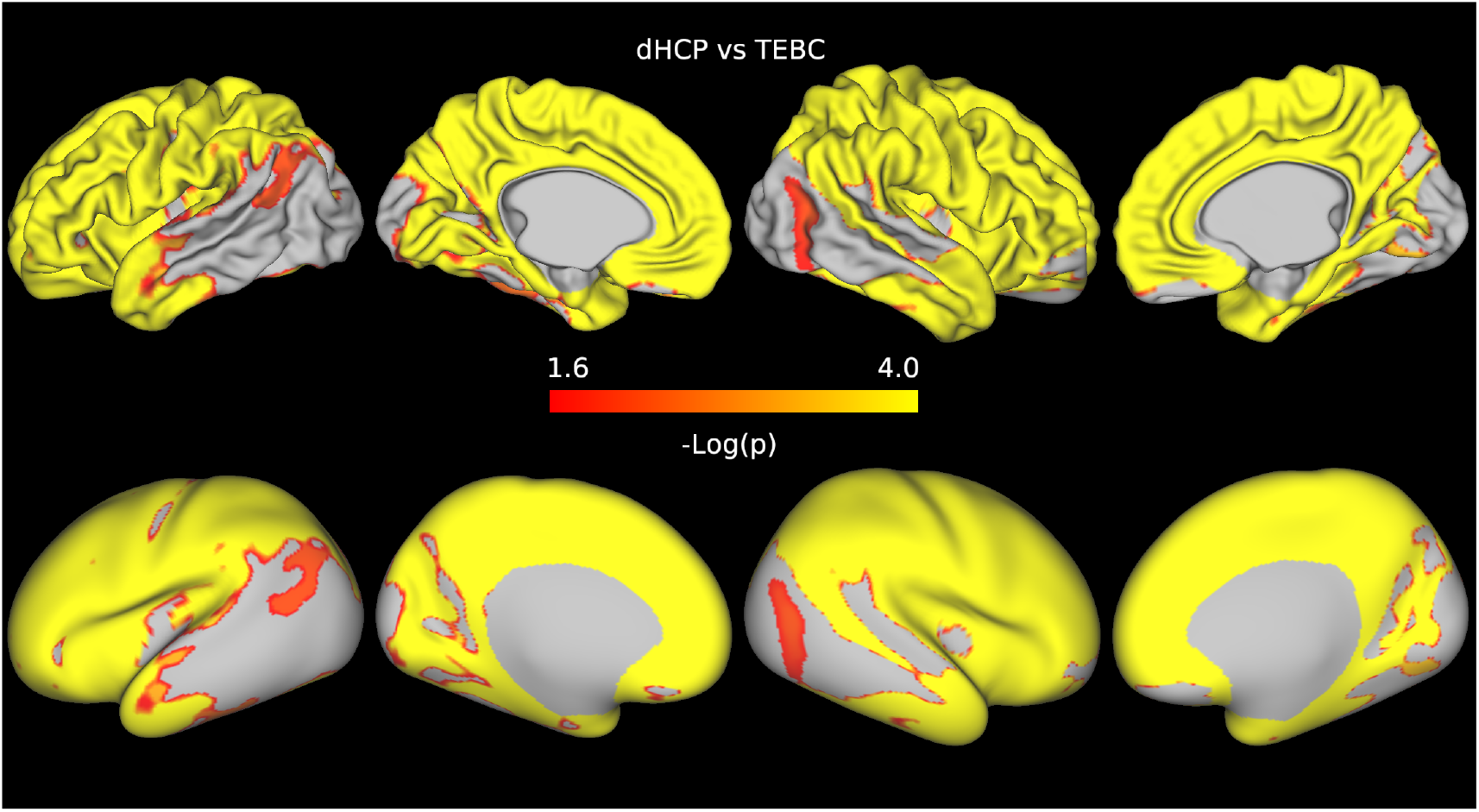
P-value maps showing the regions where the Vogt-Bailey index was higher in the TEBC sample, after controlling for prematurity, age at scan and sex.

**Figure 8.**
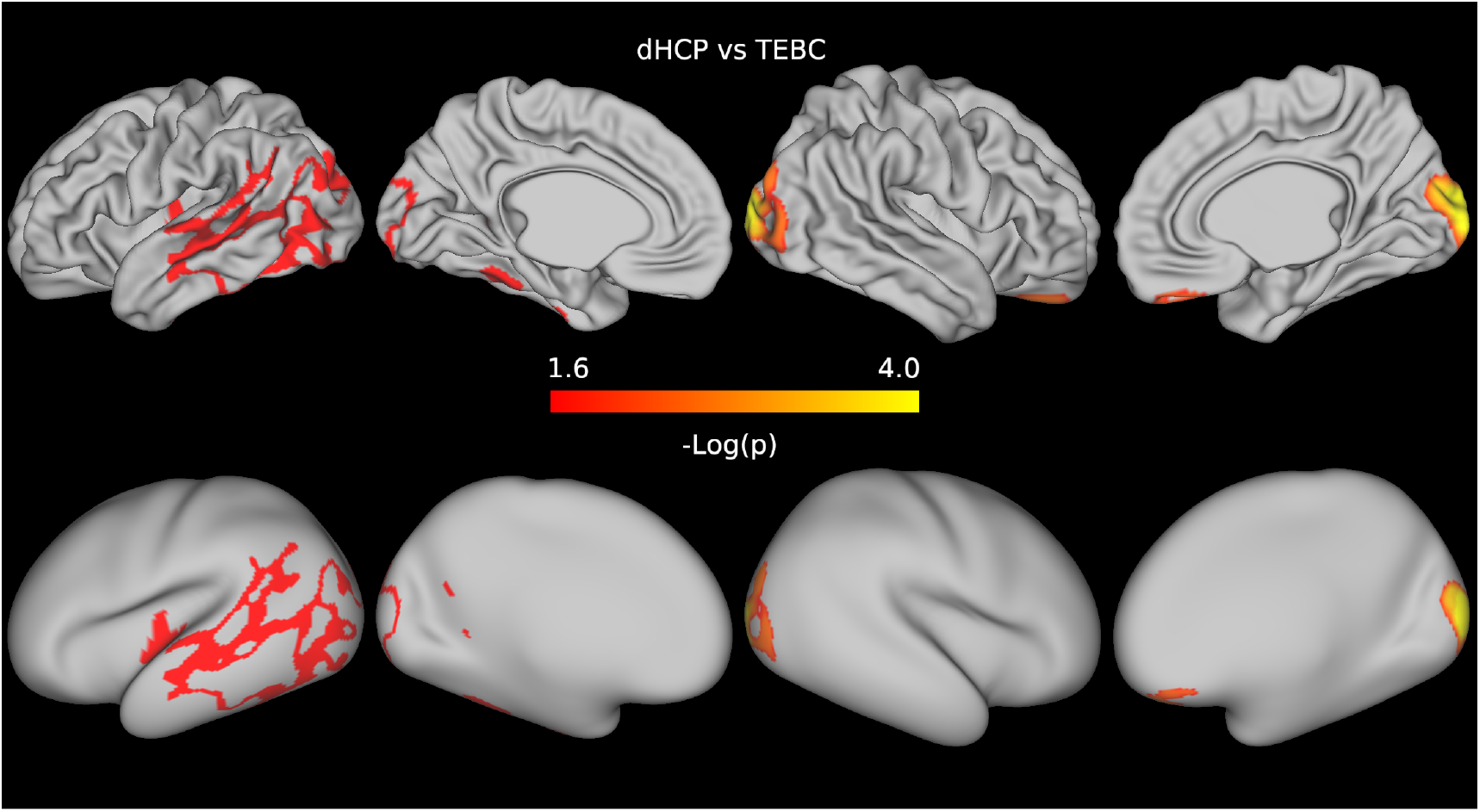
P-value maps showing the regions where the Vogt-Bailey index was higher in the dHCP sample, after controlling for prematurity, age at scan and sex.

## Discussion

In this work, we demonstrate that the VB index can be used to explore graded changes in the microstructural properties of the neonatal cortex. Consistent with previous descriptions of the dysmaturation following preterm birth as a substantially whole-brain phenomenon (***Telford et al., 2017***; ***Blesa et al., 2020***; ***Vaher et al., 2022***), across two independent datasets we find that a significant proportion of cortical microstructure is consistently altered in association with preterm birth, in a spatially distributed rather than localised fashion.

The increased homogeneity measured by the VB index in preterm infants compared to term controls could reflect increased water content, potentially as a consequence of delayed maturation, as supported by previous findings (***Ball et al., 2013b***; ***Dimitrova et al., 2021***). By interpreting the microstructural environment as a proxy for cytoarchitecture, we could speculate that in term infants local inhomogeneities are mirroring a more mature functional specialisation. In a previous study that used a region-based approach to investigate neuroanatomic variation associated with preterm birth, term infants showed less homogeneous microstructure throughout cortical gray matter regions compared to preterm infants, indicating a more advanced state of maturation (***Galdi et al., 2020***). Our results are also consistent with prior evidence that prematurity alters microstructural organisation throughout the cortex: ***Bouyssi-Kobar et al. (2018***) reported a higher diffusivity in preterm infants in the prefrontal, parietal, motor, somatosensory, and visual cortices, and ***Dimitrova et al. (2021***) reported increased cortical tissue water content and reduced neurite density index in posterior cortical regions, and greater cortical thickness in anterior cortex, in preterm compared to term-born infants. Sensory areas and posterior regions of the cortex have the largest maturational changes in microstructural and morphological profile around birth (***Garcia et al., 2018***; ***Fenchel et al., 2020***), while the network of cortico-cortical connections involving sensory-motor and transmodal integration regions is characterised by a higher complexity in term compared to preterm infants (***Blesa et al., 2021***). Together with our findings, this suggests that cortical maturation in these regions is vulnerable to the impact of preterm birth.

Previous studies have characterised the maturation of microstructure in the preterm brain by measuring deviation from developmental trajectories estimated in term subjects (***Dimitrova et al., 2020, 2021***) and have shown a high inter-individual heterogeneity in microstructure maturation throughout the cortex. Compared to these works, which examined each microstructural metric in isolation, we studied the joint variation of multiple microstructural properties to observe the impact of prematurity on cortical organisation. Our results complement these previous findings by showing that, overall, despite individual variability in deviation from normal development, prematurity results in locally homogeneous patterns of variation in a substantial portion of the cortex.

In the TEBC dataset, a region of the medial frontal lobe showed increased homogeneity of microstructure in the term subjects compared to the preterm group (Figure 5). We note that care must be taken when interpreting results along the midline sagittal plane, as they might be influenced by image processing artifacts or CSF contamination due to partial volume effects. The presence of diverging patterns of increased and decreased homogeneity could indicate that early exposure to the extra-uterine environment causes dysmaturation in a region-specific manner. Different cortical regions mature at different rates, with the insular cortex developing first, and the sensorimotor cortex maturing earlier than association cortices. This is reflected by increasing cortical anisotropy which occurs first in primary somatosensory regions and later in the frontal cortex (***Neil and Smyser, 2018***; ***Batalle et al., 2019***). Prematurity can cause deviations from typical maturational trajectories, for example with primary and non-primary auditory cortices showing a delayed maturation in preterm infants at term-equivalent age (***Monson et al., 2018***). However, we did not observe decreased homogeneity of microstructure in any region in the dHCP dataset.

The variation in results across datasets may be explained partly by unmodelled sample variability (e.g., in clinical features) and power differences, but any interpretation needs to be made in the context of the different experimental conditions. We investigated whether the observed variability was a consequence of differences in cortical thickness (that in turn might be driven by partial volume effects), and found that they could explain only in part our results. We also compared single-metric cortical maps between datasets, finding differences in most part of the cortex, but with varying spatial patterns. As our goal was to assess the replicability of VB index differences in independent datasets we did not attempt any data harmonisation, but these results serve as a caveat for future retrospective multi-site studies. State of the art harmonisation methods have the potential to minimise site-related differences especially in diffusion data (***Ning et al., 2020***), for which the processing was not harmonised in the current study. Allowing for these limitations, the existence of corresponding findings across the two datasets provides evidence of generalisability and indicates that the VB index is robust to scanner differences and methodological variation.

The VB index is a promising metric for investigating upstream determinants of cortical development. Future work could investigate whether the pattern described by the VB index underlies differences in cognitive and functional development observed in preterm infants. Elucidating the link between the widespread alterations in cortical microstructure observed in the preterm brain in the neonatal period and long-term outcomes will be important for study designs that would benefit from robust risk prediction in early life.

## Methods and Materials

In the following, we detail data acquisition and processing strategies for the two datasets. Table 4 provides a side-by-side comparison of the pipelines for structural and diffusion processing and cortical registration.

**Table 4.** Summary of acquisition parameters and processing steps for TEBC and dHCP datasets.

|  | TEBC | dHCP | Agreement |
| --- | --- | --- | --- |
| **Structural MRI** |  |  |  |
| Image Resolution | 1 mm<sup>3</sup> | 0.5 mm<sup>3</sup> | No |
| **Preprocessing** |  |  |  |
| Brain Extraction | BET | BET | Yes |
| T1 - T2 registration | Rigid Alignment, BBR | Rigid Alignment, BBR | Yes |
| Bias Correction | N4 | N4 | Yes |
| **Segmentation/Surface extraction** |  |  |  |
| Performed on | T2-weighted images | T2-weighted images | Yes |
| Tissue Segmentation | Draw-EM | Draw-EM | Yes |
| White and Pial Surface Extraction | Using tissue segmentation (no edge-based correction) | Using tissue segmentation (no edge-based correction) | Yes |
| Surface Inflation | FreeSurfer re-implementation | FreeSurfer re-implementation | Yes |
| Spherical Projection | sMDS | sMDS | Yes |
| **Diffusion MRI** |  |  |  |
| Image Resolution | 2 mm<sup>3</sup> | 1.17 * 1.17 * 1.5 mm | No |
| Bvalues (directions) | 0 (16), 200 (3), 500 (6), 750 (64), 2500 (64) | 0 (20), 400 (64), 1000 (88), 2600 (128) | No |
| **Preprocessing** |  |  |  |
| Brain Extraction | BET | BET | No |
| EPI correction | Topup | Topup | No |
| Eddy distortion correction | Eddy | Eddy | No |
| Bias correction | N4 | None | No |
| DTI fitting | dtifit | dtifit | Yes |
| NODDI fitting | Noddi-Bingham cuDIMOT | Noddi-Bingham cuDIMOT | Yes |
| **Diffusion to structural registration** |  |  |  |
| Inter modality registration | T2 to b=0 Rigid Alignment, BBR | T2 to b=1000 Rigid Alignment, BBR | No |
| **Registration to 40-week cortical template 40** |  |  |  |
| Registration to template | MSM | MSM | Yes |

BBR = Boundary-Based Registration age (*Greve and Fischl, 2009*), sMDS = spherical Multi-Dimensional Scaling (*Elad et al., 2005*), MSM = Multimodal Surface Matching (*Robinson et al., 2014*), BET = Brain Extraction Tool (*Smith, 2002*), N4 = N4 Bias Field Correction (*Tustison et al., 2010*), drawEM = draw Expectation-Maximization (*Makropoulos et al., 2014*), cuDIMOT = CUDA Diffusion Modelling Toolbox (*Hernandez-Fernandez et al., 2019*), eddy (*Andersson and Sotiropoulos, 2016*), topup (*Andersson et al., 2003*), FreeSurfer (*Fischl, 2012*), dtifit (*Jenkinson et al., 2012*).

### Data acquisition and processing

#### Theirworld Edinburgh Birth Cohort (TEBC)

Participants were recruited as part of a longitudinal study on the long-term effects of preterm birth on neurodevelopment (***Boardman et al., 2020***) between 2016 and 2021. Ethical approval was obtained from the National Research Ethics Service, South East Scotland Research Ethics Committee (11/55/0061, 13/SS/0143 and 16/SS/0154). Cohort exclusion criteria included major congenital malformations, chromosomal abnormalities, congenital infection, overt parenchymal lesions (cystic periventricular leukomalacia, haemorrhagic parenchymal infarction) or post-haemorrhagic ventricular dilatation.

A Siemens MAGNETOM Prisma 3T MRI clinical scanner (Siemens Healthcare, Erlangen, Germany) and 16-channel phased-array paediatric head receive coil were used to acquire a three-dimensional T2-weighted (T2w) SPACE (Sampling Perfection with Application-optimized Contrasts by using flip angle Evolution) structural images with 1 mm isotropic resolution, repetition time (TR) of 3.2 s and echo time (TE) of 409 ms; and a multishell axial diffusion MRI (dMRI) scan (16×b=0 s/mm^2^, 3×b=200 s/mm^2^, 6×b=500 s/mm^2^, 64×b=750 s/mm^2^, 64×b=2500 s/mm^2^) with optimal angular coverage (***Caruyer et al., 2013***). Diffusion MRI images were acquired in two separate acquisitions to reduce the time needed to re-acquire data lost to motion artifacts. In addition, an acquisition of 3 b0 volumes with an inverse phase encoding direction was performed. All dMRI images were acquired using single-shot spin-echo echo planar imaging (EPI) with 2-fold simultaneous multislice and 2-fold in-plane parallel imaging acceleration and 2 mm isotropic voxels; all three diffusion acquisitions had the same parameters (TR/TE 3.4 s/78 ms). Images affected by motion artifacts were re-acquired multiple times as required; dMRI acquisitions were repeated if signal loss was seen in 3 or more volumes. Infants were fed and swaddled and allowed to sleep naturally in the scanner. Pulse oximetry, electrocardiography and temperature were monitored. Flexible earplugs and neonatal earmuffs (MiniMuffs, Natus) were used for acoustic protection. All scans were supervised by a doctor or nurse trained in neonatal resuscitation. Full details on the protocol are provided in ***Boardman et al. (2020***).

Diffusion MRI processing was performed as follows: for each subject the first two dMRI acquisitions were concatenated and then denoised using a Marchenko-Pastur-PCA-based algorithm (***Veraart et al., 2016***; ***Tournier et al., 2019***); eddy current, head movement and EPI geometric distortions were corrected using outlier replacement and slice-to-volume registration (***Andersson et al., 2003***; ***Andersson and Sotiropoulos, 2016***; ***Andersson et al., 2016, 2017***); bias field inhomogeneity correction was performed by calculating the bias field of the mean b0 volume and applying the correction to all volumes (***Tustison et al., 2010***). Structural T2-weighted images (1 mm isotropic) were processed using the dHCP minimal processing pipeline (***Makropoulos et al., 2018***). Finally, the mean b0 EPI volume of each subject was co-registered to their structural T2w volume using boundary-based registration (***Greve and Fischl, 2009***).

#### The developing Human Connectome Project (dHCP)

Participants were scanned at the Evelina Newborn Imaging Centre, Evelina London Children’s Hospital between 2015 and 2019; the study was approved by the National Research Ethics Committee (REC: 14/Lo/1169) (***Hughes et al., 2017***). Subjects were excluded if their scans had incidental findings with possible significance for both clinical and imaging analysis (e.g. destructive white matter lesions). To build a sample that matched as closely as possible the characteristics of the TEBC dataset, we included only preterm infants that had their MRI scans acquired at term-equivalent age and we selected term controls with matching age at scan, age at birth and sex distribution using the nearest neighbour method of the R package *MatchIt* (***Ho et al., 2011***).

Preprocessed data from the third dHCP data release were used. In brief, images were acquired using a 3T Philips Achieva system (Philips Medical Systems, Best, The Netherlands). All neonates were scanned without sedation in a scanner environment optimised for neonatal imaging, including a dedicated 32-channel neonatal coil. MR-compatible ear putty and earmuffs were used to provide noise attenuation. Neonates were fed, swaddled and positioned in a vacuum jacket prior to scanning to promote natural sleep (without sedation). All scans were supervised by a neonatal nurse and/or paediatrician who monitored heart rate, oxygen saturation and temperature throughout the scan (***Hughes et al., 2017***). T2w images were obtained using a Turbo Spin Echo (TSE) sequence, acquired in two stacks of 2D slices (in sagittal and axial planes), using parameters: TR=12s, TE=156ms, SENSE factor 2.11 (axial) and 2.58 (sagittal) with overlapping slices (resolution 0.8 × 0.8 × 1.6 mm). Motion correction and super-resolution reconstruction techniques were employed resulting in isotropic volumes of resolution 0.5 mm^3^ (***Cordero-Grande et al., 2018***; ***Kuklisova-Murgasova et al., 2012***). Diffusion MRI data were acquired over a spherically optimised set of directions on three shells (b = 400, 1000 and 2600 s/mm^2^). A total of 300 volumes were acquired per subject, including 20 with b = 0 s/mm^2^. For each volume, 64 interleaved overlapping slices were acquired (in-plane resolution = 1.5 mm, thickness = 3 mm, overlap = 1.5 mm). The data were then superresolved along the slice direction to achieve isotropic resolution of 1.5 mm^3^.

Structural T2-weighted images (0.5 mm isotropic) were processed using the minimal processing pipeline of the dHCP (***Makropoulos et al., 2014, 2018***). Diffusion image preprocessing was carried out according to the dHCP diffusion processing pipeline (***Bastiani et al., 2019a***). This includes motion correction and distortion correction (***Andersson et al., 2003***; ***Andersson and Sotiropoulos, 2016***; ***Andersson et al., 2016, 2017***).

#### Cortical surface reconstruction

The surface reconstruction from T2-weighted images was carried out using a modified form of the dHCP surface processing pipeline (***Makropoulos et al., 2018***). The standard pipeline works by fitting an initial surface to boundary of the white and cortical grey matter tissue segmentation; this is subsequently refined using a novel intensity-driven force by shifting the boundary to the location of the image intensity gradient (***Schuh et al., 2017***). For this dHCP cohort this has been shown to correct segmentation errors and improve surface reconstruction but the parameters of the optimization are cohort specific and challenging to tune, often leading to failure on data with low resolution or intensity contrast. To avoid this issue and to align the processing of both datasets, all surfaces were reconstructed by running the pipeline with this feature turned off (available through version 1.1.1: https://github.com/DevelopingHCP/structural-pipeline/tree/dhcp-v1.1.1).

#### Derivation of microstructural tissue maps

DTI maps were calculated from the dMRI processed images to obtain fractional anisotropy (FA) and mean, axial and radial diffusivities (MD, AD and RD, respectively). To calculate the DTI maps, only the shells of b = 750 s/mm^2^ (TEBC) and b = 1000 s/mm^2^ (dHCP) were used. NODDI maps were calculated for both datasets using the Bingham distribution to obtain the neurite density index (NDI), orientation dispersion index (ODI) and the isotropic water fraction (ISO) (***Zhang et al., 2012***; ***Tariq et al., 2016***). The ISO map was used to obtain the tissue fraction (1-ISO), which was used to modulate the NDI and ODI maps (***Parker et al., 2021***). All diffusion metric maps were propagated to the T2w image in each subject’s native space, z-scored and concatenated to obtain for each voxel a vector comprising all six microstructural metrics.

#### Registration to a common template

Cortical surfaces were registered to the 40-week symmetric dHCP cortical template (***Williams et al., 2021***) using multimodal surface mapping (***Robinson et al., 2014, 2018***). For this, the script *align_to_template_no_volumetric_initialisation*.*sh* (https://github.com/ecr05/dHCP_template_alignment) was used. This script works by registering each subject to its age-specific template and then concatenating the subject-to-template registration with a registration which maps the age-specific template to the 40-week template (***Williams et al., 2021***). The configuration parameters were optimised for this study by including an additional regularisation level. The configuration file is available in the project repository (https://git.ecdf.ed.ac.uk/jbrl/neonatal-vbi).

#### Quality control

We used the *eddy_qc* tool (***Bastiani et al., 2019b***) to measure in-scanner motion for dMRI acquisitions. For each volume, motion is quantified by averaging voxel displacement across all voxels (computed as 3 translations and 3 rotations around the x, y and z axes). We report the average relative motion (w.r.t. the previous volume) across all volumes. Note that the motion estimates are not directly comparable across datasets due to differences in resolution and parametrization of input sequences and eddy correction. To check the quality of the registration of cortical surfaces to the template space, individual sulcal depth maps were overlaid onto template surfaces and visually inspected by three independent raters (PG, MBC and KV). Subjects were included in the analyses if five anatomical landmarks were clearly identifiable and correctly registered: central sulcus, superior temporal sulcus, lateral fissure, parieto-occipital fissure and calcarine sulcus. An example is shown in figure 9. Inter-rater agreement was 94%. The quality check was performed separately for the left and right hemispheres, resulting in two different samples, one per hemisphere.

**Figure 9.**
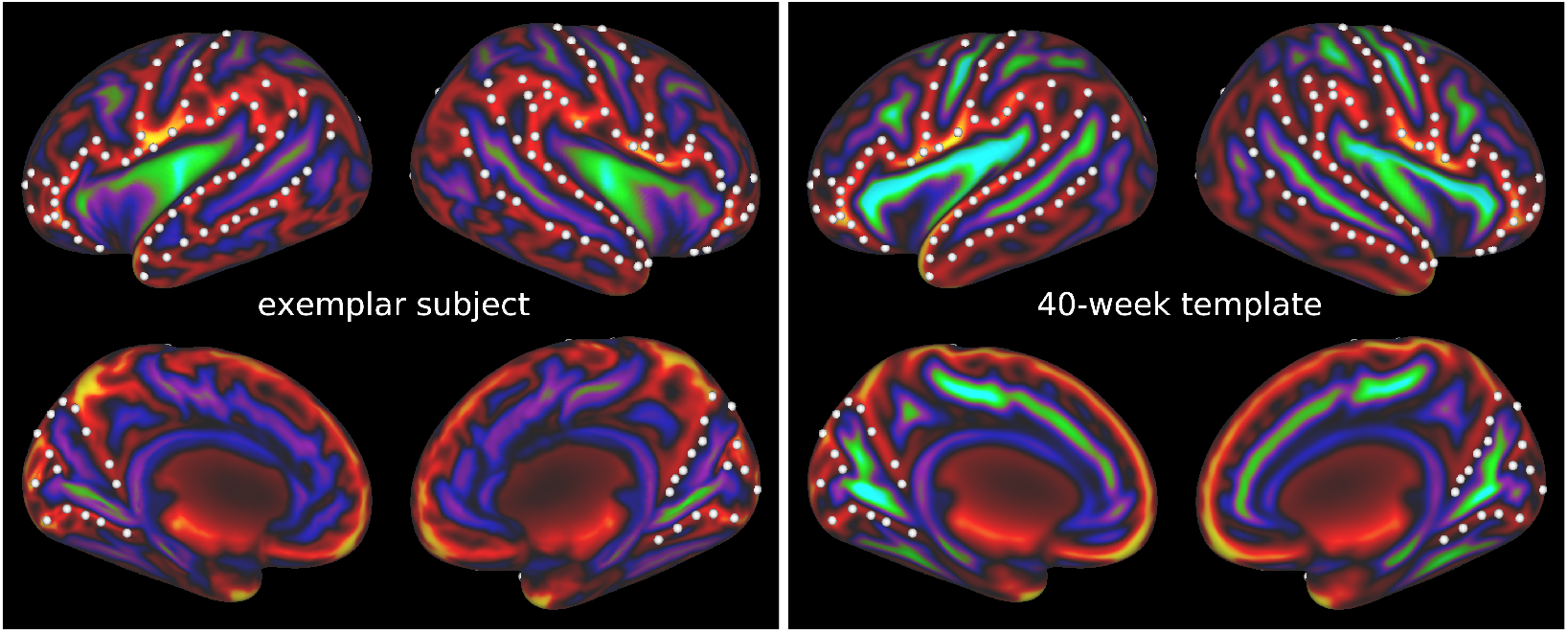
Sulcal map of an exemplar 38-week subject (left) compared to the 40-week sulcal template from dHCP (right). White dots demarcate five anatomical landmarks used for visual inspection: central sulcus, superior temporal sulcus, lateral fissure, parieto-occipital fissure and calcarine sulcus.

#### Vogt-Bailey index

Given a set of features computed at each vertex of a cortical surface, the Vogt-Bailey index measures the extent of homogeneity in cortical intra-areal relationships. Using a searchlight approach, at each vertex an affinity graph is built measuring affinity between each vertex and its neighbours; the affinity is calculated using a modified Pearson correlation ((***Bajada et al., 2020***)) computed over the features used to describe the vertices (in this case the six microstructural metrics); only positive correlations are retained. The VB index is then computed as the scaled algebraic connectivity of the affinity graph, i.e. as the second smallest eigenvalue of the Laplacian matrix of the graph. The index assumes values close to 0 when there are sharp changes in the cortical features in an area, and values close to 1 if the changes are graded. Compared to the original implementation described in ***Bajada et al. (2020***), where the neighbourhood was selected as the set of adjoining vertices on the cortical surface, in this work we used a hybrid approach where the neighbourhood is reconstructed in volumetric native space, in order to avoid spatial correlations induced by artefactual geometric patterns that follow cortical gyrification patterns (***Ciantar et al., 2022***). The coordinates of a given vertex and its neighbours are used to select the closest corresponding voxels in volumetric space, and the affinity graph is computed over the feature vectors of the matched voxels that fall within a 27-voxel cube centred on the voxel corresponding to the input vertex.

#### Statistical analysis

The *PALM* tool (***Winkler et al., 2014***) from the FSL suite (***Jenkinson et al., 2012***) was used to perform a surface-based, vertex-wise term vs. preterm comparison of the VB maps, controlling for PMA at scan and sex. An analysis comparing VB maps between datasets was also run, controlling for prematurity, PMA at scan and sex. Separate analyses were run for each hemisphere, and independently for the two datasets. Prior to the analyses, Gaussian-weighted smoothing was applied to individual VB maps, with a smoothing kernel with full-width at half-maximum set at 4 mm. Permutation p-values were computed over 10000 random shuffles with threshold-free cluster enhancement (***Smith and Nichols, 2009***) and family-wise error rate corrections. Statistical significance was set at p < 0.0253 (equivalent to *α* = 0.05, after Šidák correction over the two hemispheres, ***Šidák*** (***1967***)).

## Data availability

Requests for original TEBC image data will be considered through the BRAINS governance process (www.brainsimagebank.ac.uk). Any additional data supporting the findings of the study are available from the corresponding author, upon reasonable request. Information on how to access dHCP data is available at the project homepage (https://biomedia.github.io/dHCP-release-notes/). The cortical template and the registrations for the dHCP dataset are available at https://brain-development.org/brain-atlases/atlases-from-the-dhcp-project/cortical-surface-template/. Code to reproduce the analyses presented in this paper is available at https://git.ecdf.ed.ac.uk/jbrl/neonatal-vbi. The VB toolbox is available at https://github.com/VBIndex/py_vb_toolbox.

## Acknowledgements

We thank the families who took part in the study and the Edinburgh Imaging research staff for providing the infant scanning. This work was supported by Theirworld and was undertaken in the MRC Centre for Reproductive Health, which is funded by MRC Centre Grant [MRC G1002033]. PG was partly supported by the Wellcome Trust - University of Edinburgh Institutional Strategic Support Fund 3 [IS3-R1.1320/21]. CJB would like to acknowledge funding for the Boundaries of the Brain project (REP_2020_005) by the MCST Research Excellence Programme 2020 Call and the BE-BOB University of Malta Research Excellence Fund Grant Agreement No. 202201. MJT is supported by NHS Lothian Research and Development Office. KV is funded by the Wellcome Translational Neuroscience PhD Programme at the University of Edinburgh (108890/Z/15/Z). ECR was supported by the Academy of Medical Sciences/the British Heart Foundation/the Government Department of Business, Energy and Industrial Strategy/the Wellcome Trust Springboard Award [SBF003/1116] and a Wellcome Collaborative Award [215573/Z/19/Z]. LZJW is supported by funding from the Common-wealth Scholarship Commission, United Kingdom. The dHCP is funded by the European Research Council under the European Union’s Seventh Framework Programme (FP/2007-2013) / ERC Grant Agreement no. [319456]. For the purpose of open access, the authors have applied a Creative Commons Attribution (CC BY) license to any Author Accepted Manuscript version arising from this submission.

## Appendix 1

### Statistical comparison of cortical thickness between datasets

We compared the cortical thickness of reconstructed cortical surfaces between the TEBC and the dHCP dataset using PALM. Before running the comparison, cortical thickness maps were smoothed with a 4mm kernel. Prematurity, age at scan and sex were included as covariates of no interest. Permutation p-values were computed over 10000 random shuffles with threshold-free cluster enhancement and family-wise error rate corrections, and statistical significance was set at p < 0.0253 (equivalent to *α* = 0.05, after Šidák correction over the two hemispheres). Results are shown in figure 1. Differences were located primarily in the medial part of the cortical surfaces, including medial portion of the frontal and posterior temporal cortices, and in the parietal cortices, where cortical thickness was higher in the TEBC data. In contrast, higher thickness values in the dHCP dataset were found in the superior frontal gyrus and orbito-frontal cortex, the temporal pole and the medial occipital cortex.

**Appendix 1 Figure 1.**
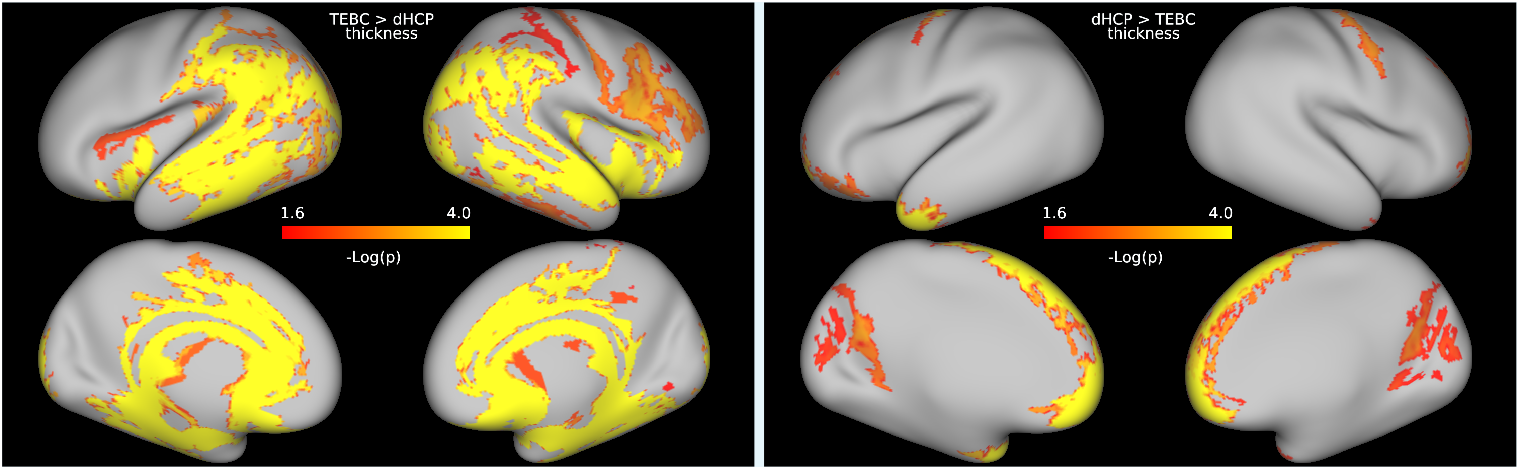
P-value maps showing the regions where cortical thickness was higher in the TEBC sample (left) and in the dHCP sample (right), after controlling for prematurity, age at scan and sex.

## Appendix 2

### Statistical comparison of microstructural maps between datasets

To further investigate the variability of results across datasets, we designed an additional experiment to compare microstructural maps between the TEBC and the dHCP dataset. To this end, subject metric maps in volumetric native space were projected onto individual cortical surfaces. The obtained surface maps were smoothed and registered to the 40-week dHCP cortical template before running a vertex-wise dataset comparison in PALM, controlling for prematurity, age at scan and sex. As for the other experiments, permutation p-values were computed over 10000 random shuffles with threshold-free cluster enhancement and familywise error rate corrections, and statistical significance was set at p < 0.0253 (equivalent to *α* = 0.05, after Šidák correction over the two hemispheres). Results are shown in figures 1 to 6. Differences were distributed throughout the cortex for all the metrics. MD, AD and RD showed similar patterns, with higher values for the TEBC dataset in the temporal and occipital lobes, and higher values for the dHCP dataset in the fronto-parietal and insular cortex. FA maps showed higher values in the TEBC dataset in the medial prefrontal, cingulate and anterior temporal cortices, while in the dHCP dataset FA values were higher in the occipital and parietal lobe and posterior temporal cortex. In the NDI and ODI maps, the TEBC dataset showed higher values in the cingulate and medial fronto-pariental cortices and the right lateral prefrontal cortex, and the dHCP dataset showed higher values in the occipital, parietal and temporal lobes and the left frontal cortex.

**Appendix 2 Figure 1.**
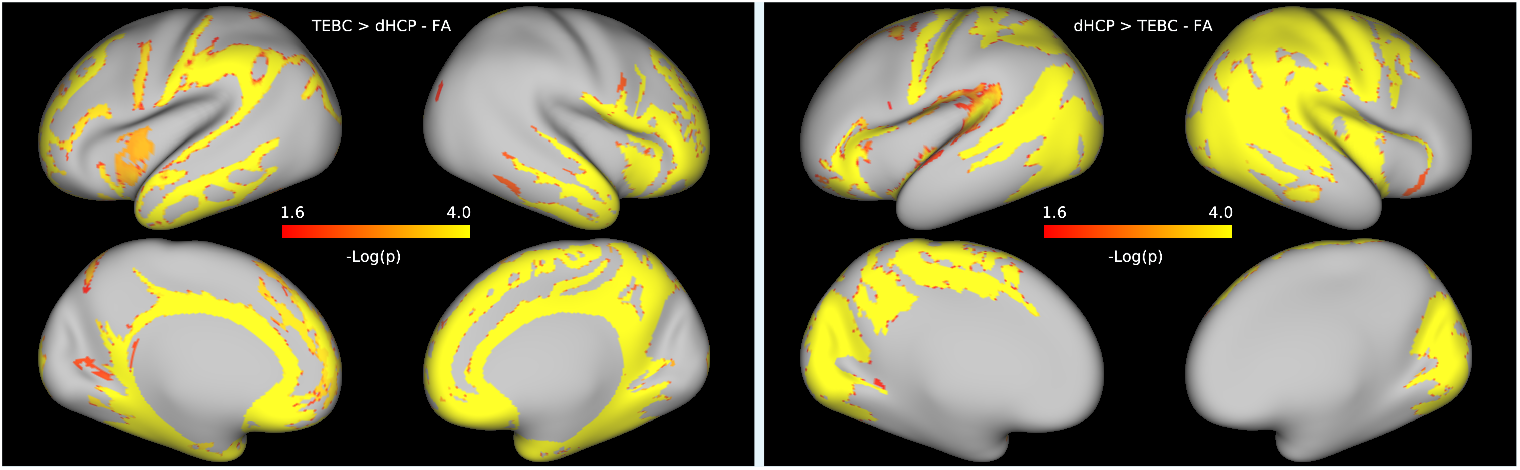
P-value maps showing the regions where FA was higher in the TEBC sample (left) and in the dHCP sample (right), after controlling for prematurity, age at scan and sex.

**Appendix 2 Figure 2.**
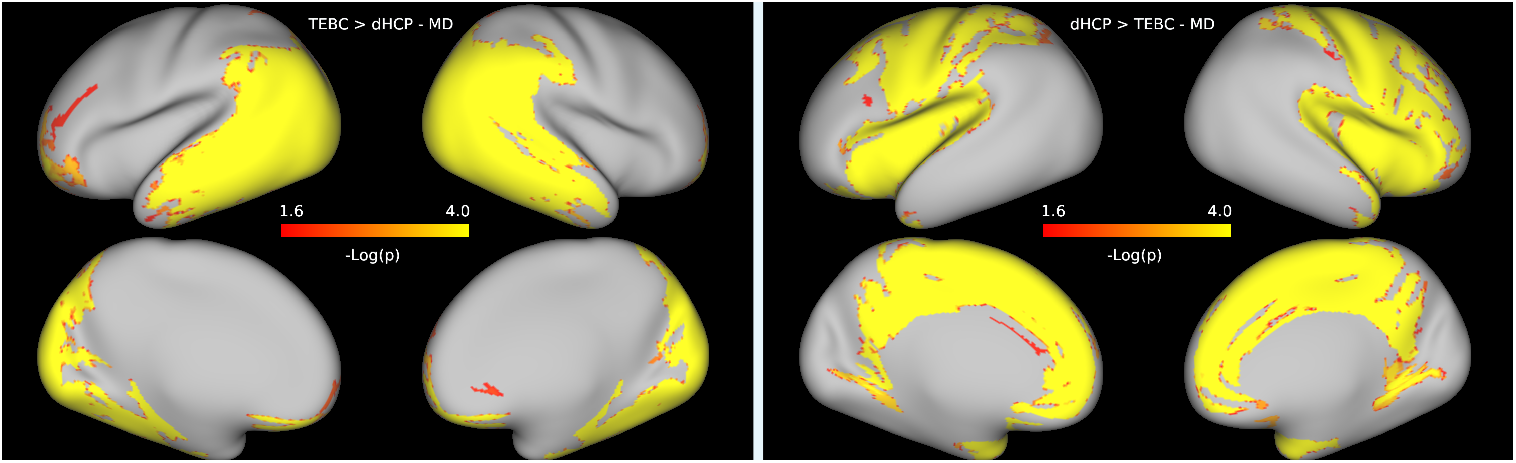
P-value maps showing the regions where MD was higher in the TEBC sample (left) and in the dHCP sample (right), after controlling for prematurity, age at scan and sex.

**Appendix 2 Figure 3.**
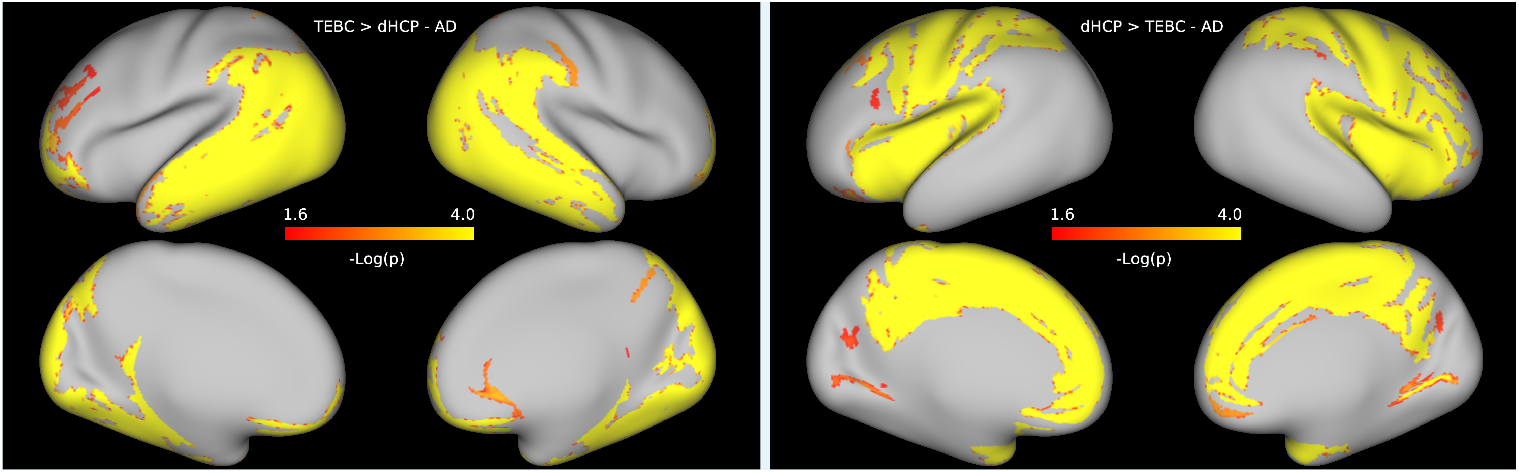
P-value maps showing the regions where AD was higher in the TEBC sample (left) and in the dHCP sample (right), after controlling for prematurity, age at scan and sex.

**Appendix 2 Figure 4.**
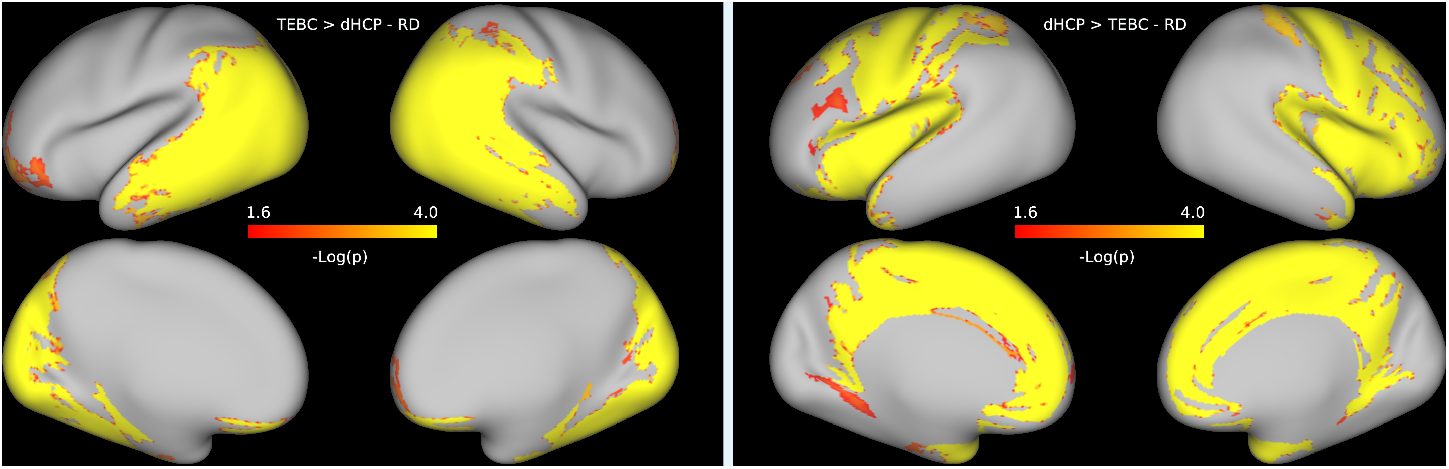
P-value maps showing the regions where RD was higher in the TEBC sample (left) and in the dHCP sample (right), after controlling for prematurity, age at scan and sex.

**Appendix 2 Figure 5.**
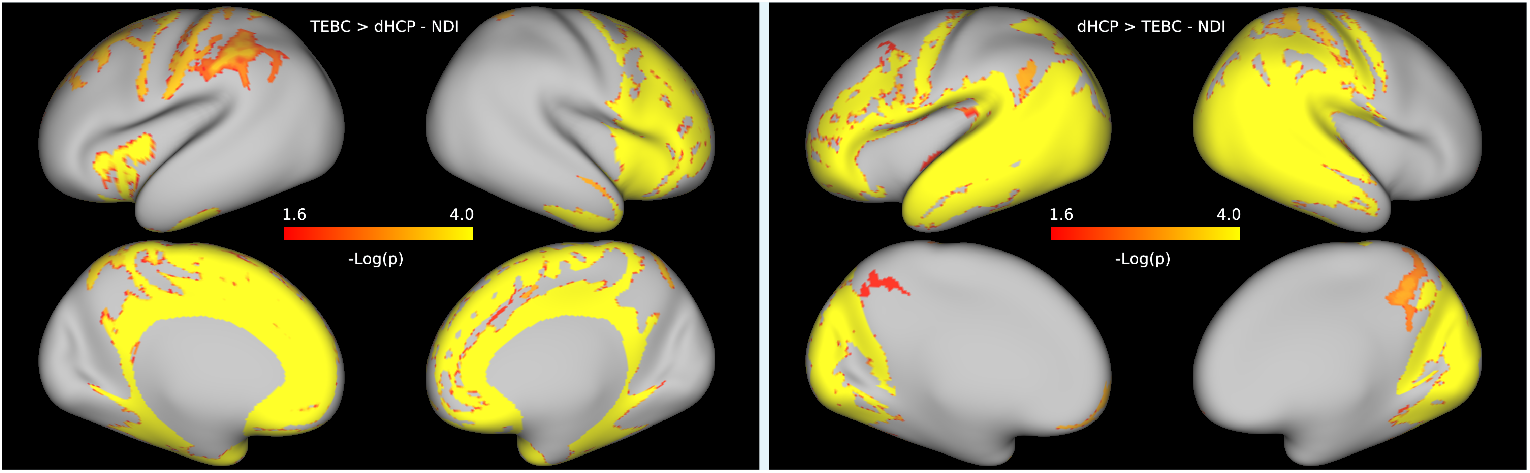
P-value maps showing the regions where NDI was higher in the TEBC sample (left) and in the dHCP sample (right), after controlling for prematurity, age at scan and sex.

**Appendix 2 Figure 6.**
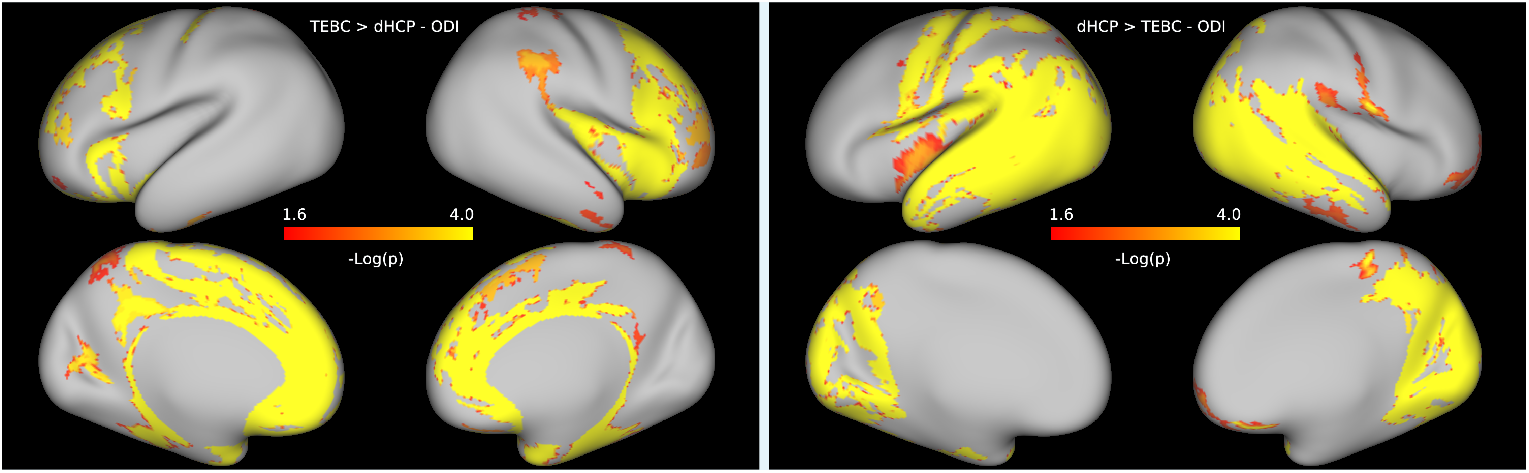
P-value maps showing the regions where ODI was higher in the TEBC sample (left) and in the dHCP sample (right), after controlling for prematurity, age at scan and sex.

## Notes

### Competing Interest Statement

The authors have declared no competing interest.

